# Local ecdysone synthesis in a wounded epithelium sustains developmental delay and promotes regeneration in *Drosophila*

**DOI:** 10.1101/2024.02.25.581888

**Authors:** Douglas Terry, Colby Schweibenz, Kenneth Moberg

**Affiliations:** Graduate Programs in Genetics and Molecular Biology, Laney Graduate School, Emory University; Graduate Programs in Biochemistry, Cell, and Developmental Biology, Laney Graduate School, Emory University; Department of Cell Biology, Emory University School of Medicine, Atlanta, GA 30322

## Abstract

Regenerative ability often declines as animals mature past embryonic and juvenile stages, suggesting that regeneration requires redirection of growth pathways that promote developmental growth. Intriguingly, the *Drosophila* larval epithelia require the hormone ecdysone (Ec) for growth but require a drop in circulating Ec levels to regenerate. Examining Ec dynamics more closely, we find that transcriptional activity of the Ec-receptor (EcR) drops in uninjured regions of wing discs, but simultaneously rises in cells around the injury-induced blastema. In parallel, blastema depletion of genes encoding Ec biosynthesis enzymes blocks EcR activity and impairs regeneration but has no effect on uninjured wings. We find that local Ec/EcR signaling is required for injury-induced pupariation delay following injury and that key regeneration regulators *upd3* and *Ets21c* respond to Ec levels. Collectively, these data indicate that injury induces a local source of Ec within the wing blastema that sustains a transcriptional signature necessary for developmental delay and tissue repair.

## INTRODUCTION

Regeneration of damaged tissue often involves cells around the injury reentering the cell cycle to repopulate the damaged area. In some cases, local stem cells are the main source of new cells while in others adjacent uninjured cells partially dedifferentiate and divide. Regenerative capacity varies widely between species and is often lost at post-embryonic or juvenile stages, suggesting it depends on redirection of developmental pathways toward regenerative growth. The associated pause in developmental growth ensures synchrony by allowing injured tissue to repair before rejoining a normal developmental trajectory.

In *Drosophila melanogaster* larvae, genetically induced death of epithelial cells in the wing pouch results in proliferation of surviving cells (the ‘blastema’) that restores adult wing size. This process is accompanied by concentration of Wingless (Wg) morphogen within the blastema, and by blastema-specific gene expression, including the transcription factor *Ets21C* and secreted factors *upd3* and *Ilp8* (*Drosophila* insulin-like peptide-8), which enhance blastema proliferation and suppress production of the steroid hormone ecdysone (Ec) in the prothoracic gland (PG) (Smith-Bolton et al., 2009, Worley et al., 2022, Romao et al., 2021, Colombani et al., 2015, Colombani et al., 2012, Garelli et al., 2012, Garelli et al., 2015). This decline in circulating Ec elicits a pause in development by reducing activity of its cognate receptor EcR. Mutations that impair key elements of this pause mechanism or that reduce injury-induced proliferation each reduce wing regeneration efficiency.

Loss of regeneration competence in a commonly used *Drosophila* wing injury system parallels a surge of Ec at the late larval-to-pupal transition (Smith-Bolton et al., 2009, Harris et al., 2016). This temporal correlation has led to the hypothesis that high Ec inhibits regeneration (Narbonne-Reveau and Maurange, 2019). However, Ec can promote regeneration in other species and is present during regeneration-competent stages of the *Drosophila* life cycle at lower levels that promote growth in the absence of injury (Das and Durica, 2013, Dye et al., 2017, Herboso et al., 2015, Karanja et al., 2022). Moreover, Ec/EcR are required for activity of the Dpp and Wg pathways (Parker and Struhl, 2020), which promote normal and regenerative wing disc growth.

The paradox that injury depletes circulating Ec at a time when the wing blastema is undergoing regeneration led us to assess Ec roles within injured discs. We find that as EcR activity drops in uninjured disc regions disc, it rises in blastema region. Local depletion of Ec biosynthesis enzymes has a limited effect on growth of uninjured wings but strongly impairs regrowth of injured wings; depleting the Ec catabolic enzyme Cyp18a1 reciprocally enhances regeneration. Overall, our data indicate that Ec/EcR signaling is required for injury-induced pupariation delay and regeneration, and that mRNAs encoding the regeneration regulators Upd3 and Ets21c are responsive to Ec. These data support a model in which the blastema is a specialized signaling environment that generates a local Ec source necessary for efficient tissue repair.

## RESULTS and DISCUSSION

### Local knockdown of Ec synthesis/degradation genes modulates wing disc regeneration

To assay ecdysone (Ec) roles during wing disc regeneration, we used an established wounding system utilizing *rotund (rn)-GAL4* and *tubulin-GAL80^ts^* transgenes to express a pulse of the apoptotic gene *reaper* in the larval pouch (hereafter *rn^ts^>rpr*) (**Fig. 1A**) (Smith-Bolton et al., 2009). Flies lacking *UAS-rpr* are used as uninjured (‘mock’; *rn+^ts^*) controls. Combining the *rn^ts^* injury system with RNAi lines is an effective approach to identify proteins within the pouch that promote regeneration (Skinner et al., 2015, Khan et al., 2017, Tian and Smith-Bolton, 2021, Brock et al., 2017, Narbonne-Reveau and Maurange, 2019). We adapted this local knockdown (KD) approach to test enzymes that synthesize Ec or its bioactive form 20-hydroxyecdysone (20E) (collectively termed *Halloween* genes) (Petryk et al., 2003, Warren et al., 2004, Chavez et al., 2000, Gilbert, 2004) (**Fig. 1B**).

**Figure 1.**
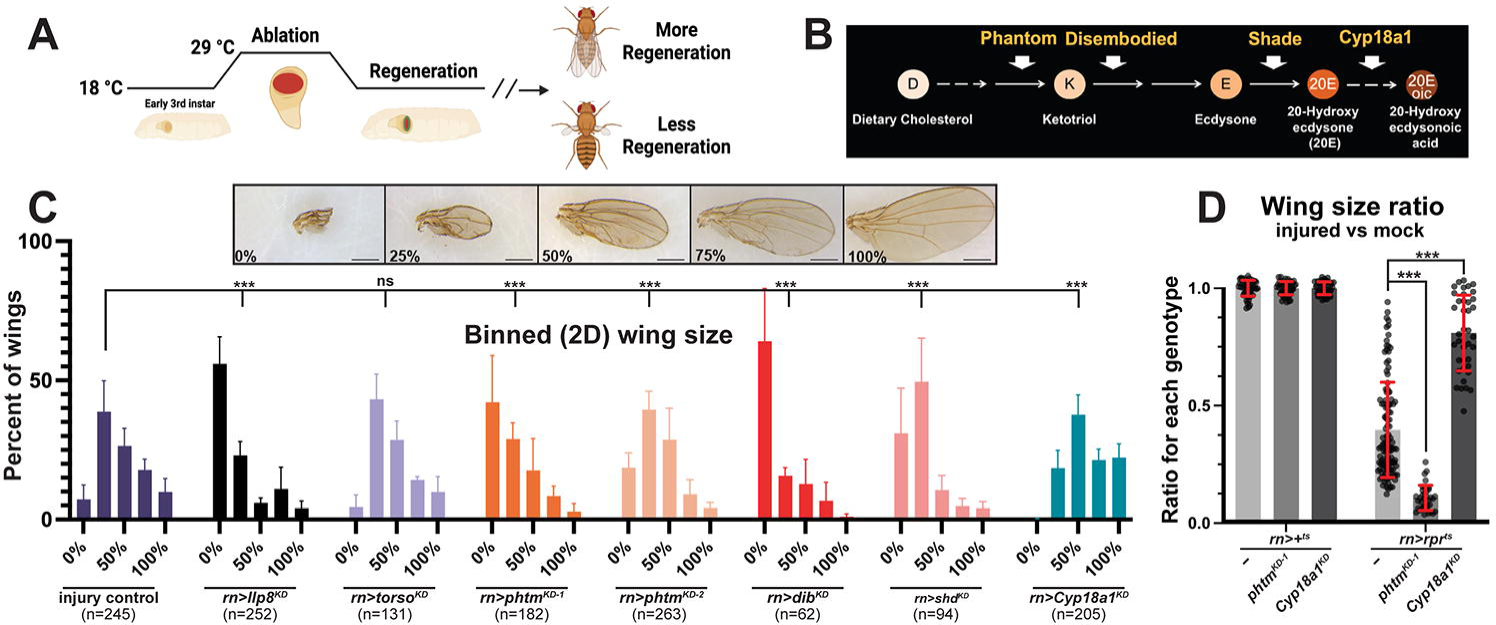
Local Ec synthesis gene knockdown impairs regeneration. **(A)** *rn>rpr^ts^* genetic ablation system. **(B)** *Halloween* pathway schematic. **(C)** Binned adult female wing sizes from 0-100% regeneration with examples. Scale bars=0.5 mm. χ^2^-test *\*\*\*p<*0.0001, Error bars=s.e.m. **(D)** Wing sizes of control (-), *phtm^KD^*, and *Cyp18a1^KD^* injured (*rn>rpr^ts^*) normalized to uninjured (*rn*>+*^ts^*) of the same genotype. *rn>rpr^ts^* n=174; *rn>rpr^ts^+pthm^KD-1^* n=100; *rn>rpr^ts^+Cyp18a1^KD^* n=78; *rn>+^ts^* n=57; *rn>phtm^KD^* n=73; *rn>Cyp18a1^KD^* n=57. Unpaired t-test: \*\*\**p*<0.0001. Error bars=s.d.

Binning adult wings by size reveals that local KD of *Halloween* genes *phantom* (*pthm*), *disembodied* (*dib*), or *shade* (*shd*) in *rn^ts^>rpr^ts^* animals significantly impair regeneration relative to *rn^ts^>rpr* alone; KD of the Ec catabolic enzyme *Cyp18a1* (Guittard et al., 2011) reciprocally enhances regeneration (**Fig. 1C**). KD of the *torso* (*tor*) receptor that promotes *Halloween* gene expression in the PG (Rewitz et al., 2009, Yamanaka et al., 2013) does not affect wing regeneration, while depletion of the peptide hormone *Ilp8* impairs regeneration, as described previously (Boone et al., 2016, Boulan et al., 2019, Colombani et al., 2015, Colombani et al., 2012, Garelli et al., 2012, Garelli et al., 2015, Gontijo and Garelli, 2018, Skinner et al., 2015). Quantification of wing area ratios (each genotype relative to its corresponding uninjured control; **Fig. 1D**) in *control* (-), *phtm^KD^*, and *Cyp18a1^KD^* confirmed that *pthm^KD^* prevents adult wings from regrowing to normal size, while *Cypa18a1^KD^* enhances regrowth. *pthm^KD^* and *Cyp18a1^KD^* each slightly reduced the absolute size of uninjured wings, perhaps due to non-specific effects of RNAi (**Fig. S1**).

### EcR is transcriptionally active in regenerating wing discs

Given the requirement for Ec biosynthesis enzymes in the regenerating wing disc, we assessed EcR activity in injured discs using *7xEcRE-GFP*, a reporter line with seven EcR response elements (EcREs) upstream of *GFP* (Hackney et al., 2007). Without injury, *7xEcRE-GFP* activity increases between early and mid-third instar (L3), elevated EcR activity detected in the hinge, pouch, and sensory organ precursors along the dorsoventral boundary (**Fig. 2A-B**). *7xEcRE-GFP* patterns are unaffected by local *pthm^KD^* in the pouch (**Fig. 2C-E**), consistent with the PG as the major Ec source in larvae.

**Figure 2.**
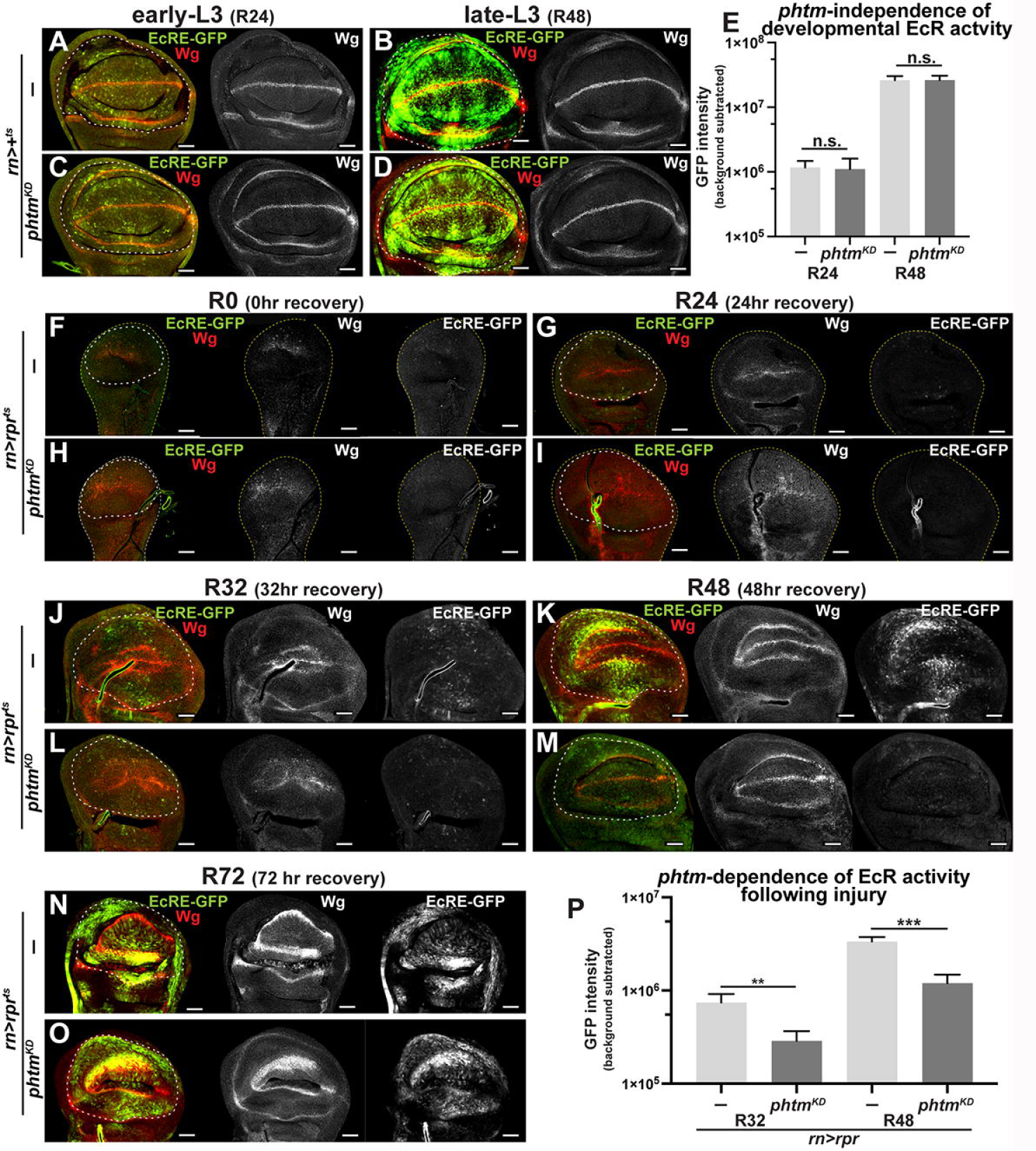
EcR is active in the blastema mid-regeneration. **(A-D)** Representative images of *EcRE-GFP* expression in mock ablated (*rn>+^ts^*) control (-) or *phtm^KD^* discs at R24 and R48 timepoints. **(E)** Quantification of *EcRE-GFP* expression in genotypes indicated in (**A-D**). R24 control; n=14, R24 *pthm^KD-2^* n=15; R48 control n=12; R48 *pthm^KD-2^* n=12. (unpaired t-test. Error bars=s.e.m.) (**F-O**) Representative images of *EcRE-GFP* activity and Wg during a R0-to-R72 time-course following *rn>rpr^ts^* injury in control and *pthm^KD-2^* discs. **(P)** Quantification of *EcRE-GFP* expression in R32 and R48 pouches (dotted lines). R32 control n=21; R32 *pthm^KD-2^* n=16; R48 control n=18, R48 *pthm^KD-2^* n=20. Unpaired t-test:\*\**p*<0.005, \*\*\**p*=0.0007, nError bars=s.e.m. **E,P** Y-axis**=**log(10). White dotted lines outline pouch/blastema as determined by Wg expression in hinge. Scale bars=50μm.

During the first 24hrs of recovery after *rpr* injury (R0-R24), *7xEcRE-GFP* expression drops to almost undetectable levels in control and *pthm^KD^* injured discs (**Fig. 2F-I**). By R32 (mid-regeneration), *7xEcRE-GFP* expression rises in scattered dorsal/ventral hinge cells and within the pouch proper (**Fig. 2J**) and by R48 is strengthened along the ventral hinge and the inner dorsal hinge cells (**Fig. 2K**), which have been shown to contribute to the wing regenerate (Smith-Bolton et al., 2009, Herrera et al., 2013, Worley et al., 2022). By late regeneration at R72, *7xEcRE-GFP* rises again and begins to resemble its pattern in uninjured mid-L3 wing discs (e.g., **Fig. 2B**). Importantly, local *pthm^KD^* in the blastema suppresses *7xEcRE-GFP* in R32 and R48 discs (**Fig. 2J-M, P**) but has little effect on *7xEcRE-GFP* in R72 discs (**Fig. 2N-O**). Quantification of *7xEcR-GFP* within the pouch and inner hinge region confirms reliance on *phtm* specifically during the R32-R48 period (graphs in **Fig. 2P** and **Fig. S2O**; see **Fig. S2A-N** for a R0-R72 timecourse), implying a transient reliance on local Ec midway through regeneration.

Use of the TNF receptor ligand *eiger* (*rn>egr^ts^*) as an alternative to *rpr* strongly induces *7xEcRE-GFP* and *EcRE-lacZ* in the pouch and decreases in the hinge and notum, indicating that elevated EcR activity at the site of injury and a drop elsewhere are also features of the *egr* injury system (**Fig. S2P-S**). Notably, *7xEcRE-GFP* expressing cells in *rn>egr^ts^* injured discs overlap the Wg-positive blastema (**Fig. S2T**), express high levels of EcR (**Fig. S2U**), and are adjacent Dcp-1 (caspase) positive cells (**Fig. S2V**). *rn>egr^ts^* regeneration is also impaired by *shd^KD^* or the *shd^2^* null allele (**Fig. S2W**). For subsequent experiments, we used the *rn>rpr^ts^* system due to a direct apoptotic mechanism and less crumpled disc morphology (Smith-Bolton et al., 2009, Harris et al., 2016, Igaki et al., 2002, Igaki et al., 2009, Yoo et al., 2002).

### Injury-induced pupariation delay is regulated by *pthm* and *Cyp18a1* in the wing pouch

Previous studies have shown that Upd3 and Ilp8 secreted from growth-perturbed wing discs trigger developmental delay by suppressing systemic production of Ec in the PG (Smith-Bolton et al., 2009, Worley et al., 2022, Romao et al., 2021, Colombani et al., 2015, Colombani et al., 2012, Garelli et al., 2012, Garelli et al., 2015), which delays the larval-to-pupal transition and allows injured wing discs to regrow and equalize with uninjured discs prior to pupation. Removing Ilp8 impairs this delay, and thus prevents efficient regeneration of adult wings (Hackney et al., 2012, Halme et al., 2010, Jaszczak et al., 2016, Colombani et al., 2012, Garelli et al., 2012, Skinner et al., 2015). *pthm^KD^* or *Cyp18a1^KD^*. Significantly, *pthm^KD^* suppresses injury-induced pupariation delay to a similar extent as an *Ilp8^KD^* control, while *Cypa18a1^KD^* has the opposite effect of enhancing the delay; local *torso^KD^* did not affect pupation timing (**Fig. 3A-B**). All RNAi lines tested slightly accelerated pupariation (**Fig. S3A-B**) and slightly reduce wing size (**Fig. S1**), which could suggest non-specific effects of the RNAi processing machinery.

**Figure 3.**
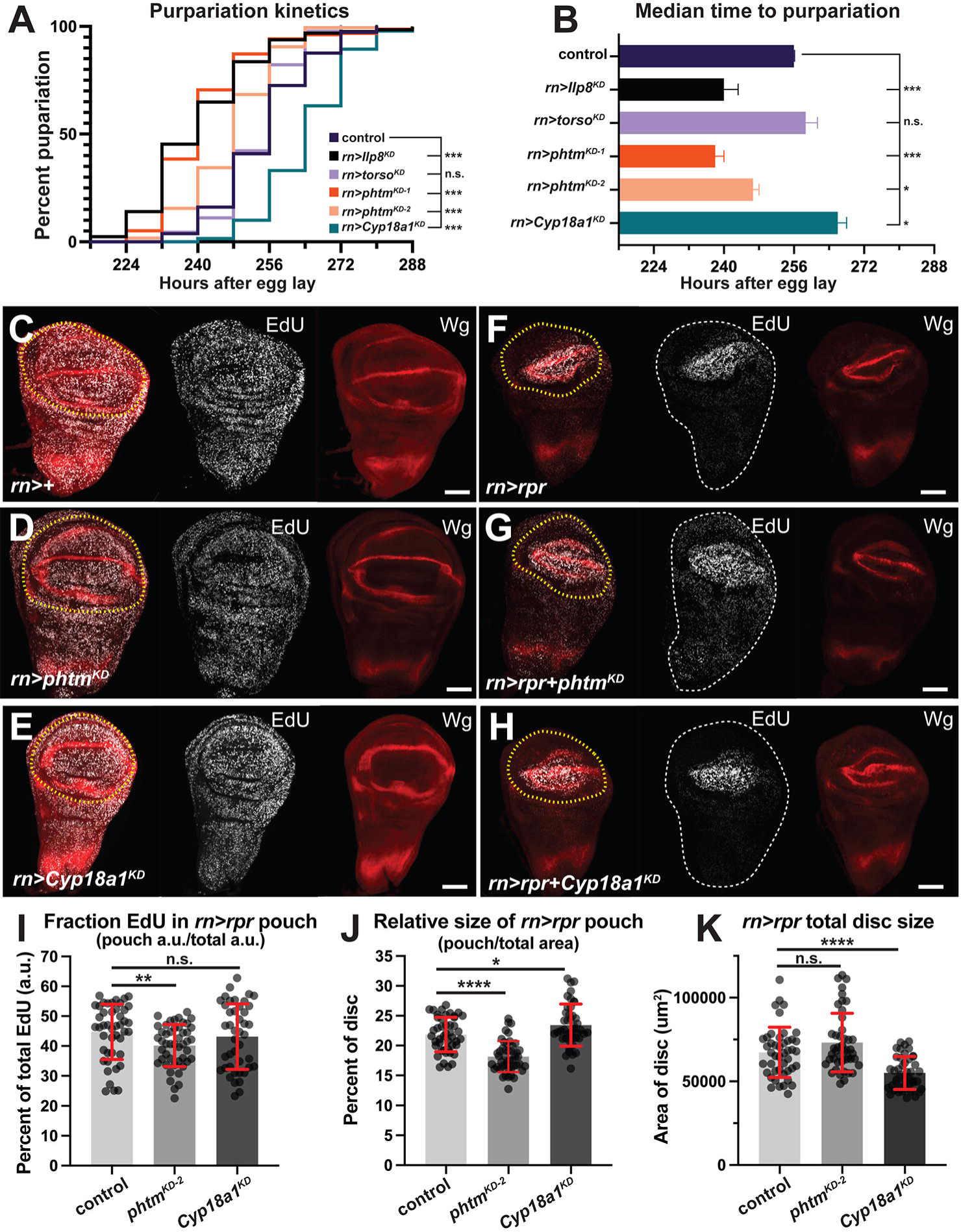
Injury induced pupariation delay and coordinated intra-organ growth is dependent on local Ec. **(A)** Kaplan-Meyer curve of pupariation kinetics following injury (*rn>rpr^ts^*). Log-rank (Mantel-Cox) test; control n=186; *pthm^KD-1^* n=156; *pthm^KD-2^* n=180; *Cyp18a1^KD^* n=190; *torso^KD^* n=130; *ilp8^KD^* n=128. **(B)** Median time to pupariation following injury of genotypes in (**A)** (One-way Anova, Dunnett’s post-hoc test). **(C-H)** Representative images of EdU i and Wg in R48 (**C-E**) mock ablated (*rn*>+) control, *pthm^KD-2^*, and *Cyp18a1^KD^* discs, or (**F-H**) injured (*rn>rpr^ts^*) control, *pthm^KD-2^*, or *Cyp18a1^KD^*. Yellow dotted lines indicate pouch/blastema region used in quantifications. **(I)** EdU fluorescence and **(J)** blastema size as a percentage of disc total (blastema/disc) at R48 for indicated genotypes. **(K)** Total area of *rn>rpr^ts^* discs at R48. Control n=45; *pthm^KD-2^* n=47; *Cyp18a1^KD^* n=43. Unpaired t-test. \*\*\*\**p*<0.00001, ***p<0.0001, \*\**p*=0.0086, \**p*<0.05. Error bars=s.d. Scale bars 50μm.

### Coordinated intra-organ growth reduction is local Ec dependent

Given the pro-growth Ec role in discs (Dye et al., 2017, Herboso et al., 2015, Karanja et al., 2022, Gokhale et al., 2016, Nogueira Alves et al., 2022), we assessed effects of *pthm^KD^* and *Cyp18a1^KD^* on cell division within the blastema, focusing on R48 when *7xEcRE-GFP* activity is most dependent on *pthm* (see **Fig. 2**). In injured discs, *pthm^KD^* does not significantly alter the absolute level of pouch EdU relative to injured control (-) (**Fig. S3C**) but does reduce the fraction of EdU located with the pouch relative to overall total EdU (**Figs. 3F-H, I**). This effect seems attributable to a partial reduction of EdU incorporation in areas outside the pouch of *pthm^KD^* injured discs (**Fig. 3F-H**; see dotted white outline of whole disc area). Slowed cell division and reduced EdU in regions outside the blastema is a well-documented feature of injury-induced growth perturbation and results from inhibition of Ec production in the PG, which systemically slows growth of uninjured tissue (Boulan et al., 2019, Hackney et al., 2012, Halme et al., 2010, Jaszczak et al., 2016, Skinner et al., 2015, Worley et al., 2022, Smith-Bolton et al., 2009, Kiehle and Schubiger, 1985). Consistent with a local role for Ec and EcR in this intra-organ feedback mechanism, *pthm^KD^* has a regional effect on growth; the blastema of *pthm^KD^* discs comprises a smaller percentage of the total disc area relative to injured controls, whereas the blastema of *Cyp18a1^KD^* discs is enlarged relative to total disc area (**Fig. 3J**) and exhibits an upward trend in the fraction of EdU signal within the pouch region (**Fig. 3H-I**). *pthm^KD^* discs are larger than injured controls, and *Cyp18a1^KD^* discs are smaller than injured controls at the same time point in recovery (**Fig. 3K**). In mock ablated discs, EdU fluorescence within the pouch is not significantly changed by *phtm^KD^* and is slightly decreased by *Cyp18a1^KD^* (**Fig. S3E**). The EdU pouch fraction (pouch EdU fluorescence/total EdU fluorescence) is unchanged in uninjured *phtm^KD^* and *Cyp18a1^KD^* discs relative to control (-) (**Fig. 3C-E** and **S3F**), which mirrors the very mild effect of *pthm^KD^* or *Cyp18a1^KD^* wing size (see **Figs. 1D** and **S1**). Parallel analysis of mock-injured *rn>+* discs did not detect significant differences in overall disc or pouch size (**Fig. S3G-H**). Parallel analysis of the mitosis (M-phase) marker phospho-Histone H3 (pH3) detects a drop in mitotic index of injured pouch cells depleted of *phtm* or *dib* relative to injury alone (*rn>rpr^ts^*) (**Fig. S3I-T**), suggesting that local Ec is also required for maximal mitotic activity within the blastema. Overall, these data are consistent with *phtm* impairing intra-organ signaling required to inhibit growth of disc regions outside the pouch, and with a potential role for *phtm* in promoting mitotic progression.

### The regeneration genes *upd3* and *Ets21C* respond to Ec

Based on a prior RNA-seq study that identified *phtm* mRNA among a group of injury-induced transcripts in sorted cells from *rn>rpr^ts^* blastemas (Khan et al., 2017), we reanalyzed a public single-cell RNA sequencing (scSeq) data derived from injured (*rn>egr^ts^* system) and uninjured larval wing discs (Floc’hlay et al., 2023). This data confirmed *phtm* induction in wound cells and further refined this expression to ‘beta’ wound cells (**Fig. S4A-B, H)** (as defined in Floc’hlay et al., 2023). Expression of *sro* (*shroud*), a *Halloween* gene that acts upstream of *phtm* in the Ec synthesis pathway, is detected by scSeq throughout the disc independent of injury (**Fig. S4C**). Following injury, the Ec-inducible genes *ImpE1*, *ImpL2*, *ImpL3* (Paine-Saunders et al., 1990, Moore et al., 1990, Osterbur et al., 1988, Natzle et al., 1988, Natzle et al., 1986) and *Cyp18a1*, which is induced by EcR as part of an feedback inhibitory loop (Rewitz et al., 2010) are induced in blastema cells (**Fig. S4D-H)**. Reanalysis of public CUT&RUN data (Uyehara and McKay, 2019) indicate that all four of these loci can be bound by EcR in larval wing cells (**Fig. 4SI**).

Based on evidence of Ec synthesis and EcR activation in injured wings, we assessed effects of *pthm^KD^*, *dib^KD^*, or *Cyp18a1^KD^* on mRNAs encoding *upd3*, *Ilp8*, and the regeneration-specific transcription factor *Ets21c* (**Fig. 4**) (Worley et al., 2022, Santabarbara-Ruiz et al., 2015, Katsuyama et al., 2015, Pastor-Pareja et al., 2008, La Fortezza et al., 2016). Pouch-specific depletion of *pthm* or *dib* in the background of *rn>rpr^ts^* significantly impairs injury-induction of *upd3* mRNA relative to uninjured controls (as detected by qPCR) at R48 (48hrs recovery), while depletion of *Cyp81a1* increases *pthm* expression at R48 (**Fig. 4A**). A *upd3-lacZ* enhancer trap reports elevated *upd3* transcription in the ventral hinge at R24 of *rn>rpr^ts^* discs (**Fig. 4B, left panels**), which overlaps the region of *phtm*-dependent EcR activity post-injury (see **Fig. 2**). By R48, *upd3-lacZ* appears in large, apically located pouch cells that resemble the pattern of EcR protein detected in post-injury discs (**Fig. S2U**). Importantly, R24 and R48 *upd3-lacZ* expression are reduced when tested in age-matched *rn>rpr^ts^*+*phtm^KD^* disc (**Fig. 4B**, **right panels** and **Fig. S5**). Quantification of fluorescence intensity confirms dependence of *upd3-lacZ* on local *phtm* (**Fig. 4C**). We find no evidence that *phtm*, *dib*, or *Cyp18a1* have roles in expression of *Ilp8* mRNA or a *Ilp8-GFP* reporter in injured (**Fig. 4A,D**) or uninjured discs (**Fig. S6A**). *phtm^KD^* and *dib^KD^* also have no significant effect on *Ets21* levels; however, *Cyp18a1^KD^* strongly elevates *Ets21c* mRNA levels at both time points post-injury despite the observation that (**Fig. 4A**). This *Cyp18a1* effect may suggest that excess Ec can promote *Ets21c* mRNA accumulation but is not normally required for it. Recent evidence indicates that wing injury alters epithelial permeability (DaCrema et al., 2021) and could thus affect the ability of Upd3 and Ilp8 to escape the disc lumen and signal to distant tissues. We measured this parameter using fluorescein conjugated dextran (FITC-dextran) but found no evidence that this parameter responds to local *pthm^KD^* or *Cyp18a1^KD^* (**Fig. S6B)**.

**Figure 4.**
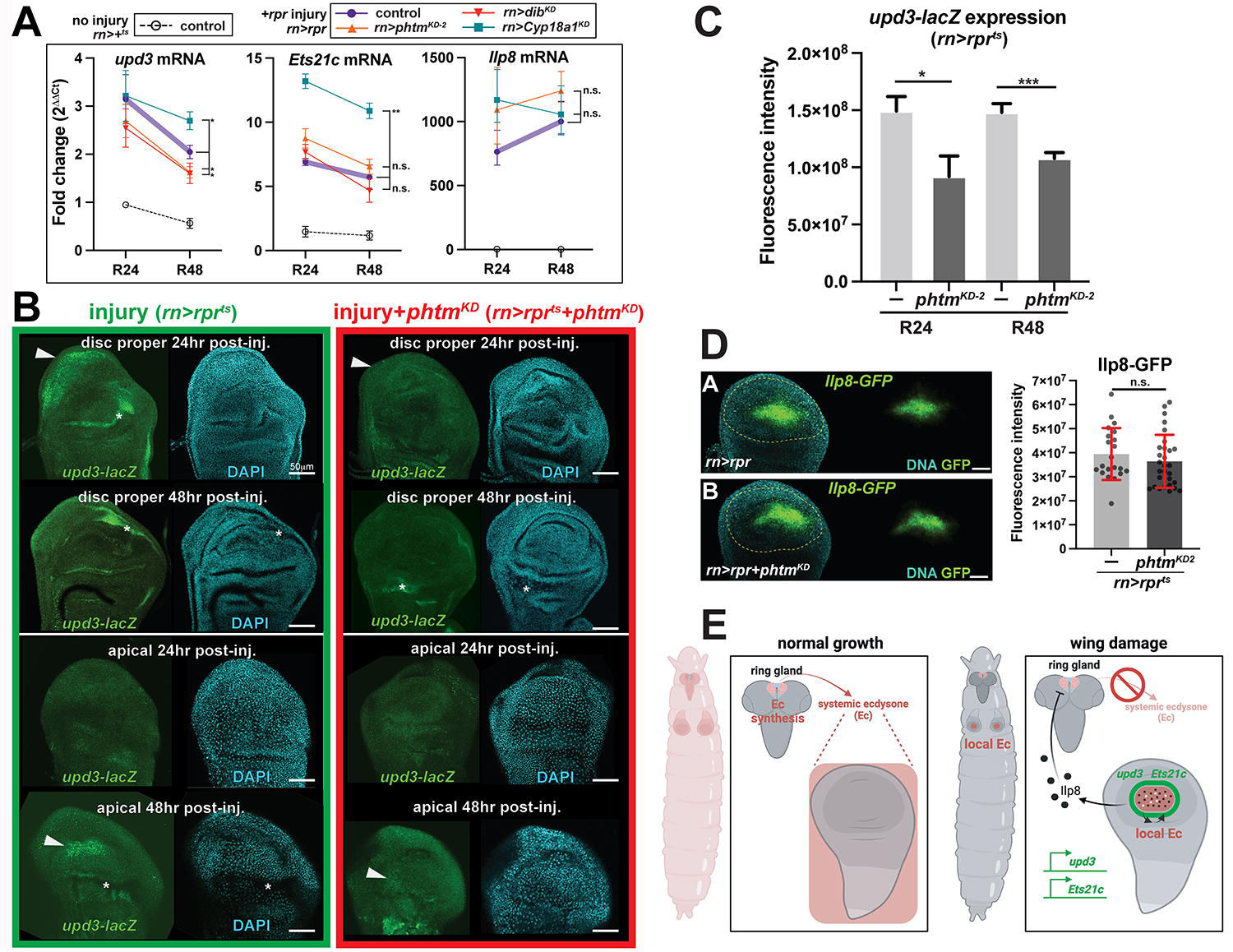
*Upd3* and *Ets21C* levels, but not *Ilp8* levels, are dependent on local Ec. **(A)** Fold-change in levels *Ilp8*, *Ets21c*, or *upd3* mRNA levels (injured/mock) at R24 and R48 for *rn>rpr^ts^* with *pthm^KD-2^*, *dib^KD^*, or *Cyp18a1^KD^*. Significance values for R48 data. (Unpaired t-test: \*\**p*=0.0085, \**p*<0.05. Error bars=s.e.m.). **(B)** *upd3-lacZ* expression in injured (*rn>rpr^ts^*) control (green frame) or *pthm^KD-2^* (red frame) discs at R24 or R48. Arrowheads indicate *upd3-lacZ* expression in disc proper or apical cells. Asterisks indicate non-specific staining in disc folds. Scale bars=50μm. **(C)** Quantification of *upd3-lacZ* in z-stack projections of the indicated ages (R24 or R48) and genotypes. (R24 control n=6, R24 *pthm^KD-2^* n=6; R48 control n=18, R48 *pthm^KD-2^* n=18. Unpaired t-test. Error bars=s.e.m. \*\*\**p*<0.0001, \**p*<0.05.). **(D)** Representative images of *Ilp8-GFP* expression post-injury (R48) +/- *phtm^KD^*, and quantification of *Ilp8-GFP* intensity using unpaired t-test. *Ilp8-GFP* n=21 and *pthm^KD-2^*+*Ilp8-GFP* n=29. Error bars=s.d., scale bars=50μm. **(E)** Model depicting Ec sources in the presence and absence of *rn>rpr* injury. Without injury, Ec from the PG provides a temporal cue to coordinate developmental transcriptional programs across larval tissues. Following *rn>rpr* injury, Ec production is inhibited in the PG but increases in the blastema, where it is required for local EcR activity and maximal *upd3* expression, and potentially to maintain blastema-specific regeneration factor *Ets21c*.

Based on these collective findings, we propose that local Ec influences wing disc regeneration by enhancing local expression *upd3* and potentially *Ets21c* (**Fig. 4E**). This enhanced dependence on a local Ec in genetically wounded discs contrasts with cells in developing wing discs that normally respond to Ec derived from the prothoracic gland (Parker and Struhl, 2020). As EcR activity and nuclear accumulation of EcR protein itself remains restricted to the pouch and hinge of *rn>rpr^ts^* or *egr^ts^* damaged discs (e.g., see **Figs. 2J-L** and **S2U**), locally available Ec appears to signal over a relatively short range. It is worth noting that this paradigm of local Ec synthesis driving local EcR activity is similar to the initiation of border cell (BC) migration in the ovary (Domanitskaya et al., 2014), which relies on generated Ec generated by a few follicle cells to promote invasion of the BC cluster between nurse cells (Bai et al., 2000). How Ec is concentrated locally in these two scenarios is unclear, but could rely on secretion, uptake, or local sequestration, perhaps including low levels of Ec produced by the PG during the developmental pause. EcR activity has also been shown to be induced within larval wing discs damaged by exposure to gamma irradiation (Karanja et al., 2022), which parallels the effect of *rn>rpr^ts^* and *egr^ts^* observed in this study. However, the source of Ec following irradiation was not examined. Intriguingly, activating EcR in irradiated discs by Ec feeding, or blocking EcR using a dominant-negative transgene (*EcR^DN^*), each affect expression of the *Ilp8-GFP* reporter and Wg protein, which plays a key role in regeneration (Smith-Bolton et al., 2009). Neither of these factors respond to knockdown of *Cyp18a1* or *Halloween* genes in our *rn>rpr^ts^* damaged wing discs, implying potential differences in effects of Rpr-killing vs. ionizing radiation, or in methods used to identify EcR candidate targets. Broadly, the identity of gene targets that respond to the pulses of EcR activity detected in the pouch of *rn>rpr^ts^* and *egr^ts^* models would reveal much about how a damaged wing disc evades the systemic loss of Ec from the PG by upregulating *Halloween* gene activity in the pouch, and how this local Ec contributes to the unique transcriptional signature necessary for tissue repair.

## Acknowledgments

We thank Developmental Studies Hybridoma Bank (DSHB), Bloomington Drosophila Stock Center (BDSC), Harvard Transgenic RNAi Project (TRiP), and Vienna Drosophila Resource Collection (VDRC) for antibodies and stocks. Model figure created with BioRender.com. *7xEcRE-GFP* was a gift of V. Henrich. Ablation stocks: *rn-GAL4, tub-GAL80ts*; *rn-GAL4, tub-GAL80ts, UAS-rpr*; and *rn-GAL4, tub-GAL80ts* were gifts of Rachel Smith-Bolton. We thank members of the Moberg laboratory for their discussions. Technical support from and support Emory Integrated Cellular Imaging Core Facility.Funding to D.T. from NIH F31CA239563 and to K.M. from NIH GM123136 and GM121967.

## Author contributions

Conceptualization, D.T. and K.M.; Methodology, D.T. and K.M.; Investigation, D.T., and C.S.; Writing – Original Draft, D.T. and K.M.; Writing – Review & Editing, D.T. and K.M.; Supervision, K.M.; Funding Acquisition, D.T. and K.M.

## MATERIALS AND METHODS

### Drosophila strains

The *Drosophila* lines *rn-GAL4, tub-GAL80^ts^*; *rn-GAL4, tub-GAL80^ts^, UAS-rpr*; and *rn-GAL4, tub-GAL80^ts^, UAS-egr* were previously described (Smith-Bolton et al., 2009; gifts from R. Smith-Bolton), as was *7xEcRE-GFP* (Hackney et al., 2007; gift from V. Henrich) and *upd3-lacZ* ((gift of I. Hariharan Bunker et al., 2015) The following lines were obtained from the Bloomington *Drosophila* Stock Center: *Ilp8-GFP* (*Mi[MIC]Ilp8*^MI00727^) (BDSC# 33079), *UAS-torso-RNAi* (BDSC# 58312), *UAS-phm-RNAi* (BDSC# 55392), *UAS-shd-RNAi* BDSC #67356) (Jugder et al., 2021), *shd^2^* (BDSC# 4219), *UAS-cyp18a1-RNAi* (BDSC# 64923) (Idda et al., 2020), and *7xEcRE-lacZ* (BDSC# 4517). The following lines were obtained from the Vienna *Drosophila* Stock Center: *UAS-dilp8-RNAi* (v102604) (Kashio et al., 2016), *UAS-phm-RNAi* (v104028) (Beebe et al., 2015), *UAS-dib-RNAi* (v101117) (Deady et al., 2015), *UAS-shd-RNAi* (v108911) (Knapp and Sun, 2017).

### Genetic ablation experiments

Ablation experiments were performed as previously described (Smith-Bolton et al., 2009). We used the (1) *w^1118^;;rn-GAL4, tub-GAL80ts,UAS-reaper* and (2) *w^1118^;;rn-GAL4, tub-GAL80ts, UAS-eiger* genetic ablation systems to study regeneration, and (3) *w^1118^;;rn-GAL4, tub-GAL80ts* as our mock ablation control. These are abbreviated as (1) *rn^ts^>rpr*, (2) *rn^ts^>egr*, and (3) *rn^ts^>+*. Development was synchronized by collecting eggs on grape plates. Eggs were collected for 4 hours at 25°C then placed at 18°C. After 2 days at 18°C, 50 L1 larvae were picked and placed into churned fly food vials. For *rn^ts^>rpr* experiments, on day 7 after egg lay (AEL) temperature shifts were performed to induce ablation. Vials were moved from 18°C to a 29°C circulating water bath for 20 hours. For *rn^ts^>egr*, temperature shifts were performed at day 8 AEL for 40 hours. For wing disc experiments, dissections were performed at several time points during regeneration: immediately following ablation (R0), and after 24, 48, and 72 h of regeneration. Mock ablation dissections were performed at R24 or R48 after mock ablation and developmental stage (early L3-late L3) was further confirmed by colorimetric assay (Hailstock et al., 2023).

### Adult wing imaging and area measurement

Adult wings were plucked and mounted in Gary’s Magic Mountant (Canada balsam dissolved in methyl salicylate). Images were taken on a Nikon SMZ800N microscope at 25X magnification (2.5x objective x 10x eyepiece) using a Leica MC170HD camera and a Nii-LED High intensity LED illuminator. Area of wing was measured using FIJI.

### Immunohistochemistry

Larval discs were dissected in 1xPBS, fixed 20min in 4% paraformaldehyde at room temperature, rinsed 3x in 1xPBS, then permeabilized for 30min at room temperature (RT) in 1xPBS+0.3% Triton X-100 (1xPBST 0.3%). Samples were then blocked for 1 hour in 1xPBS+10% normal goat serum (NGS), and then incubated in primary antibody 1xPBST 0.1% + 10% NGS overnight at 4°C. The next day samples were washed for 5min 5x in 1xPBST 0.1%, incubated for 45min at RT in secondary antibody in 1xPBST 0.1% + 10% NGS, and washed for 5min 5x in 1xPBST 0.1%. Nuclei were then stained with DAPI/Hoescht for 10min and washed 2x in 1xPBS, before mounting discs in *n*-propyl gallate (4% w/v in glycerol). Wing discs were imaged on a Nikon A1R HD25 confocal system using 20x and 40x objectives.Primary antibodies: anti-Wg antibody (DSHB, 4D4; 1:50), anti-β-galactosidase (DSHB, 40-1a; 1:250), anti-EcR common (DSHB; 1:100), anti-cleaved DCP-1 (Cell Signaling; 1:100), and anti-Phospho-histone H3 (Cell Signaling; 1:100). Fluorescently labeled secondary antibodies were from Jackson Immunoresearch Laboratories (1:50). Nucleic acid staining was performed by incubating discs for 10 min with Hoescht 3342 (Thermo Fisher Scientific) or DAPI (Invitrogen).

### EdU assay

EdU staining was performed according to the Click-iT EdU Cell Proliferation Kit manufacturer’s instructions (Thermo Fisher Scientific; C10340). Briefly, live discs were incubated in Schneider’s medium (Thermo Fisher Scientific; 662249) with EdU at 100 μM concentration for 20min at RT in the dark. After incubation, discs were fixed in 4% PFA for 20min before proceeding with standard antibody staining as described above. Click-iT EdU detection was then performed according to the Click-iT EdU proliferation Kit manufacturer’s instructions.

### Pupariation timing experiments

Pupariation rates were quantified by counting newly formed pupae every 8 h. Timing was reported as hours after egg lay (AEL), starting at the beginning of the 4-hour egg lay period. Pupariation was determined as when animals stopped moving and darken in color.

### Quantitative reverse-transcription PCR (qPCR)

50 discs from equivalently staged larvae of each genotype were dissected and added to 100 μL of Trizol. RNA was extracted using the Qiagen RNeasy kit (Qiagen; 74106) according to manufacturer’s instructions. cDNA conversion was performed using SuperScript-III RT kit (Thermo Fisher; 18080093) according to manufacturer’s instructions using 1 μg of extracted RNA. qPCR analysis was performed in triplicate using SYBR green master mix (Qiagen, 204143) on a QuantStudio6 Real-Time PCR System. The entire assay was repeated 3 times using discs from separate ablation experiments. Refer to Table S1 for a full list of primers used. Quantitative qPCR analysis was performed using the ΔΔC_T_ method and expression levels were normalized to *rp49*. Statistical analysis was performed in GraphPad Prsim9 using the Student’s t-test.

### Dextran assay

The dextran assay was performed as previously described (DaCrema et al., 2021). Larvae were dissected, inverted, and cleaned in Schneider’s medium at the time points noted. Carcasses were then placed in 25mg/mL 10kD fluorescein-conjugated dextran (Thermo Fisher, D1820) diluted 1:8 in Schneider’s medium and incubated for 30min at room temperature in the dark while rocking. Carcasses were then washed 1x with Schneider’s medium and the fixed with 4% paraformaldehyde for 20min, before proceeding with normal antibody staining as described above.

### UMAP analysis and EcR genome association peaks in wing cells

scSeq data were analyzed using the online portal https://scope.aertslab.or/#/WingAtlas/welcome. The session loom *multiome_scRNA.scenic1x.scope*, located under sequential *wing disc*, *atlas*, and *full_multiome* tabs, was loaded and analyzed using the *Compare* feature with setting *number of displays:2*. *REWE37570709B79* (Rotund GAL4 GAL80ts Ctrl) and *REW51C2157Eeed7* (Rotund GAL4 GAL80ts Eiger) were loaded into adjacent windows with display settings *Log transform* (off), *CPM normalize* (on), *Expression based plotting* (on), *Dissociate viewers* (off). Pouch and wound annotations were added to UMAPs using built-in *broad annotation*s setting. CPMs for individual genes were annotated using the dropdown tabs and downloaded as *.csv* files for importation into Prism to generate Bubble plots. For visualization of EcR CUT&RUN peaks, data from Uyehara, McKay 2019 was downloaded from the NCBI SRA and mapped to *Dm6* genome using Galaxy tools (usegalaxy.org) and visualized using IGV viewer (igv.org). SRA files analyzed: *SRX15238797, SRX15238795, SRX5173551, SRX5173550, SRX5173549, SRX5173548*.

## Statistical analysis

### Adult wing size

Only female animals were used for these experiments, as previous research has indicated that female and male flies display statistically different regenerative capacity. Populations from separate ablation experiments cannot be meaningfully compared to one another as studies from other labs have indicated seasonal variation regeneration.

#### Semi-quantitative

for binning experiments, adult wing size was scored following injury and regeneration into 5 categories (0%, 25%, 50%, 75%, and 100%). Results were calculated per population, with n’s representing data from at least 3 separate ablation experiments. Statistical analysis was performed in GraphPad Prism9 using the χ^2^ test.

#### Quantitative

Results were calculated per population, with n’s representing data from at least 2 separate ablation experiments. For injured wing size measurements, wings were normalized to the mean wing size of mock ablation wings from the same genotype. This was done to control for the small but statistically significant difference between mock ablation controls. Statistical analysis was performed in GraphPad Prsim9 using the Student’s t-test.

### Fluorescence intensity quantification

ImageJ/FIJI software was used to quantify fluorescence intensity. All quantified images used the same imaging parameters within a given experiment. EcRE-GFP intensity was quantified by measuring the fluorescence within the wing disc pouch as defined by the outer edge of hinge Wingless (Wg) expression. To control for background variations across discs, a 52×52 pixel non-regenerating region in the notum was used to determine background EcRE-GFP expression. Statistical analysis was performed in GraphPad Prism9 using the Student’s t-test. Total n# represents 3 separate ablation experiments. Ilp8-GFP intensity was quantified by measuring the total GFP fluorescence from the Ilp8-GFP positive region of the disc using summed z-stacks. Statistical analysis was performed in GraphPad Prism9 using the Student’s t-test. Total n# represents 3 separate ablation experiments. Intensity of ßGal expressed from *upd3-lacZ* was quantified by measuring the fluorescence within the wing disc pouch as defined by the outer edge of hinge Wingless (Wg) expression from summed z-stacks through the disc. Statistical analysis was performed in GraphPad Prism9 using the Student’s t-test. Total n# represents 3 separate ablation experiments.

### EdU assay quantification

ImageJ/FIJI software was used to quantify fluorescence intensity. All quantified images used the same imaging parameters within a given experiment. EdU intensity was measured from summed z-stacks of discs. For both intensity and area measurements, the pouch region of the disc was determined by the outer edge of hinge Wg expression. Percent of total EdU fluorescence in pouch was determined: pouch EdU intensity/total disc EdU intensity. This allowed for normalization of any variation in incorporation between discs. Area of pouch was determined: area of pouch/total disc area. Statistical analysis was performed in GraphPad Prism9 using the Student’s t-test.

### pH3+ nuclei quantification

Z-stacks of 20 planes were taken through wing discs from the indicated genotypes. ImageJ/FIJI software for automatic cell counting was used to count the number of pH3+ cells. Regions of the disc were determined by morphology and Wg antibody staining.

### Pupariation timing analysis

Survival curve analysis for pupariation timing experiments was performed in GraphPad Prism9 using the Log-rank (Mantel-Cox) test. Results are calculated per population, with n’s representing data from at least 3 separate ablation experiments.

