## Supplemental Figs, Legends and Table S1 for "Local ecdysone synthesis in a wounded epithelium sustains developmental delay and promotes regeneration in *Drosophila*"

### Supplemental Figures and Legends

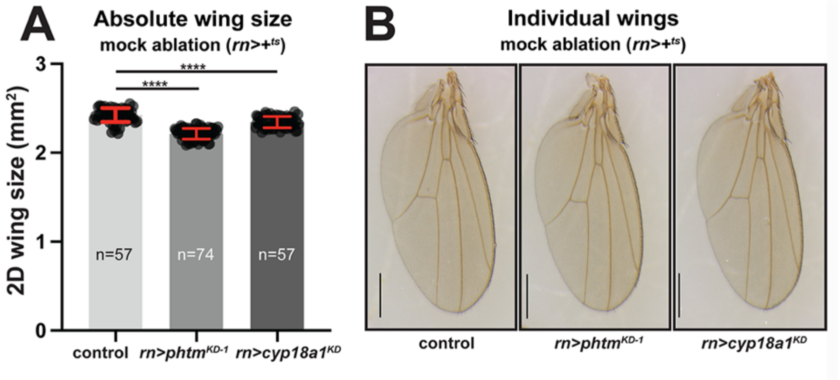


**Figure S1. Local Ec knockdown in mock ablation.**
**(A)** Absolute wing area data for mock ablated (*rn*>+) control, *pthm*$^{KD-1}$, and *Cyp18a1*$^{KD}$. Samples are female wings only. *n* values are cumulative from three separate crosses. **(B)** Representative mock ablated wings. Significance determined by Student's t-test. ****$p$<0.00001. Scale bars=0.5mm. Error bars=s.e.m.

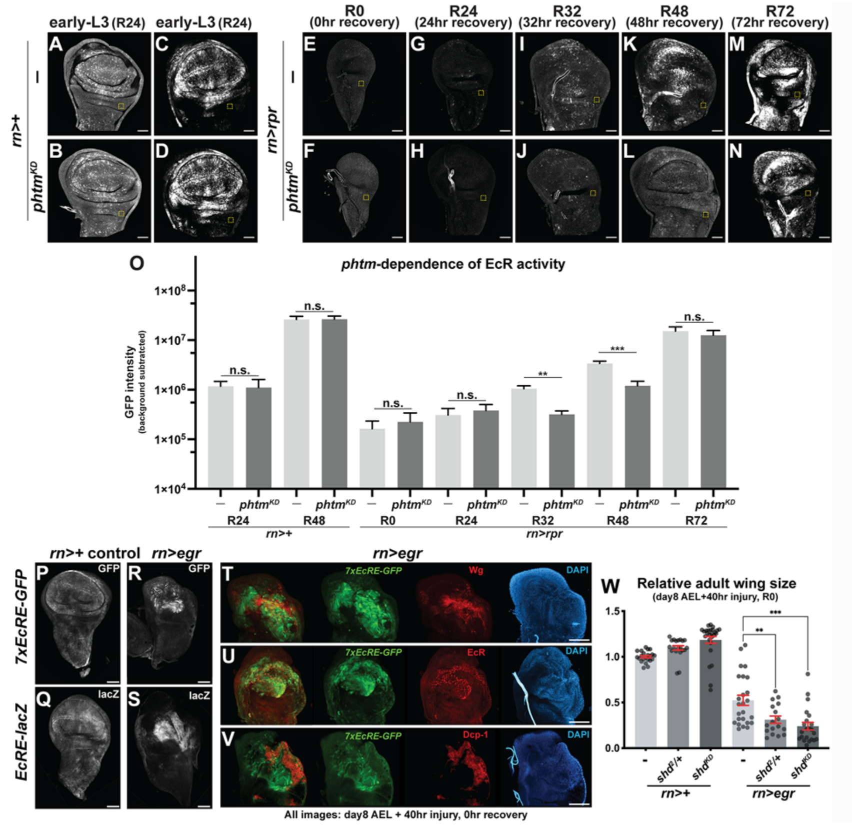


**Figure S2. Levels of EcR activity across a time course of *rn>rpr* regeneration and *shade*-dependence of EcR activity induced by the *rn>egr$^{ts}$* system.**

(**A-N**) Representative images of *7xEcRE-GFP* expression patterns in wing discs without (**A,C,E,G,I,K,M**) or with (**B,D,F,H,J,L,N**) *phtm$^{KD}$* at the indicated time points post-heat shock. **(O)** Quantification of *7xEcRE-GFP* expression in mock injured *control* discs at the *phtm*-sensitive R24-R48 timepoints compared to regenerating (*rn>rpr$^{ts}$*) discs throughout a R0-R72 time course. **mock R24**: control n=14, *pthm$^{KD-2}$* n=15; **mock R48**: control n=12, *pthm$^{KD-2}$* n=12; **injured R0**: control n=9, *pthm$^{KD-2}$* n=9; **injured R24**: control n=9, *pthm$^{KD-2}$* n=9; **injured R32**: control n=21, *pthm$^{KD-2}$* n=16; **injured R48**: control n=18, *pthm$^{KD-2}$* n=20; **injured R72**: control n=19, *pthm$^{KD-2}$* n=13. Data collected in three independent experiments. Y-axis in **(O)** is log(10) scale of fluorescence (arbitrary units). Significance determined by Student's t-test ***p*<.005, ****p*=0.0007, *n.s.*=not significant. Error bars=s.e.m. **(B-E)** Representative images of *7xEcRE-GFP* and *EcRE-lacZ* expression patterns in mock (*rn>+$^{ts}$*) and Eiger-injured (*rn>egr$^{ts}$*) discs. Scale bars=50µm. **(F-H)** *rn>egr$^{ts}$* induced expression of the *7xEcRE-GFP* reporter (green) imaged with antibodies to (**F**) Wg (red), (**G**) EcR (red; all isoforms), and (**H**) cleaved Dcp1 caspase. **(I)** Effect of *shd$^{2}$*/+ or *shd$^{KD}$* on wing size of *control* (*rn>egr$^{ts}$*) or Eiger-injured wings (*rn>egr$^{ts}$*). Sample sizes: injured (*rn>egr$^{ts}$*) n=25; *shd$^{2}$*/+ injured n=17; *shd$^{KD-2}$* injured n=22. Sizes were wings normalized to mock ablated without *shd* allele. Samples are female wings only. Student's t-test. ***p*<0.001, ****p*<0.0001. Error bars=s.e.m. Scale bars=50µm. For *rn>egr$^{ts}$* wounding, larvae were temperature shifted at Day 8 for 40h and dissected for analysis at 0R.

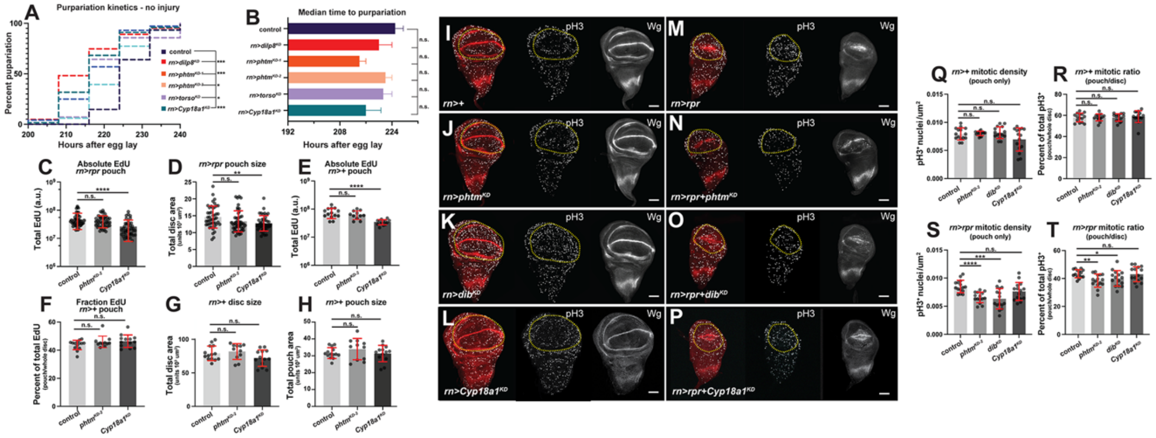


**Figure S3. Pupariation timing of uninjured larvae and analysis of EdU and mitotic index following Rpr injury.**

**(A)** Kaplan-Meyer survival curve for pupariation time-points following mock ablation (*rn*>+) with significance determined by the Log-rank (Mantel-Cox) test. Data are the sum of three independent experiments: *control* n=136, *pthm*$^{KD-1}$ n=158; *pthm*$^{KD-2}$ n=237; *Cyp18a1*$^{KD}$ n=163; *torso*$^{KD}$ n=98; *ilp8*$^{KD}$ n=184. **(B)** Median time to pupariation following mock ablation (*rn*>+) of the genotypes in (**A**). One-way Anova with Dunnett's test used to determine statistical significance. ****p*<0.0001, **p*<0.05, *n.s.*=not significant. **(C)** Total internal EdU pouch fluorescence (a.u. log[10]) and (**D**) two-dimensional (2D) pouch size from injured (*rn>rpr*$^{ts}$) wing discs expressing *phtm*$^{KD-2}$ or *Cyp18a1*$^{KD}$. Data from three separate experiments: control n=45, *pthm*$^{KD-2}$ n=47, *Cyp18a1*$^{KD}$ n=43. Student's t-test: *n.s.*=not significant. Error bars=s.d. Yellow dotted lines indicating the region of quantification based on anti-Wg staining (red) for this and following panels. (**E**) Total internal EdU fluorescence (a.u. log[10]) in mock injured (*rn*>+$^{ts}$) with the indicated RNAi lines. (**F**) Pouch fraction of EdU, (**G**) overall disc size, and (**H**) pouch size of mock *control* (*rn*>+$^{ts}$) mock ablated discs expressing *phtm*$^{KD-2}$ or *Cyp18a1*$^{KD}$. Data gathered in three separate experiments: control n=18, *pthm*$^{KD-2}$ n=16, *Cyp18a1*$^{KD}$ n=16. Student's t-test: *****p*<0.0001, ***p*<.005, n.s.=not significant. Error bars=s.d. (**I-P**) Images of anti-pH3 staining (greyscale) in **(I-L)** control (*rn*>+$^{ts}$) discs or (**M-P**) *rn>rpr*$^{ts}$ discs in combination with *pthm*$^{KD-2}$, *dib*$^{KD}$, or *Cyp18a1*$^{KD}$. Data were gathered in three independent experiments: control n=22, *pthm*$^{KD-2}$ n=27, *dib*$^{KD}$ n=18, *Cyp18a1*$^{KD}$ n=16. Student's t-test: n.s.=not significant. Error bars=s.d. **(M-P)** Representative images of pH3$^{+}$ staining in control, *pthm*$^{KD-2}$, *dib*$^{KD}$, and *Cyp18a1*$^{KD}$ ablated (*rn>rpr*$^{ts}$*)* wing discs. **(Q,S)** Mitotic density (pH3$^{+}$ nuclei/pouch μm$^{2}$) and (**R,T**) mitotic ratio (pH3$^{+}$ pouch/pH3$^{+}$ total) of mock injured (*rn*>+$^{ts}$) or injured (*rn>rpr*$^{ts}$) discs with or without *phtm*$^{KD-2}$ or *Cyp18a1*$^{KD}$. Data gathered in three seperate experiments: *control*, n=41; *pthm*$^{KD-2}$, n=37; *dib*$^{KD}$, n=44; *Cyp18a1*$^{KD}$, n=24. Student's t-test: *****p*<0.0001, ****p*=0.0005, ***p*<.005, **p*<0.05, n.s.=not significant. Error bars=s.d. Scale bars=50μm.

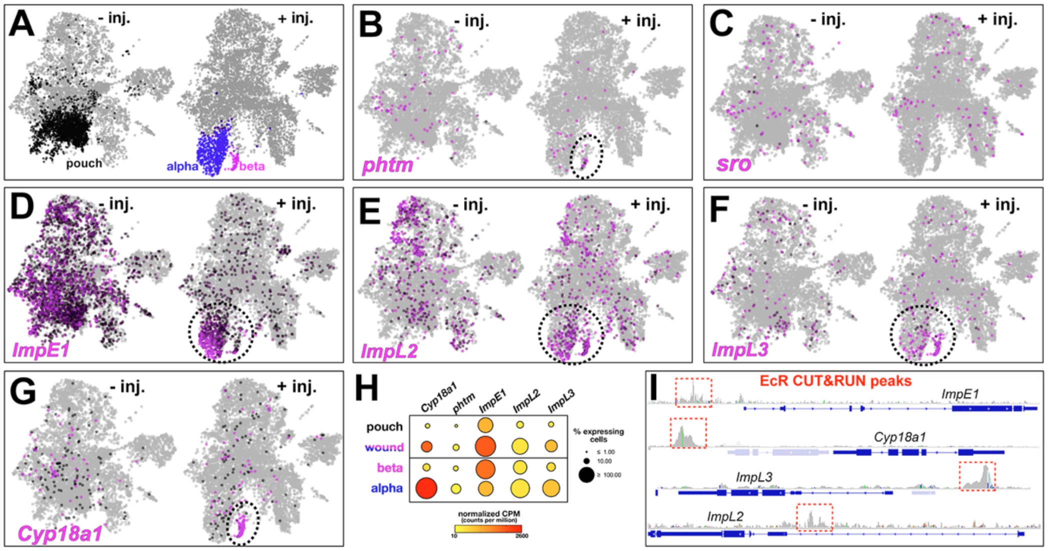


**Figure S4. Ilp8-GFP levels in mock ablation and epithelial barrier permeability in *pthm*$^{KD}$.** UMAP plots of larval wing discs generated from a public single cell sequencing database (https://scope.aertslab.org/#/WingAtlas/WingAtlas) with annotations for (**A**) pouch (black), and alpha wound (blue) or beta wound (magenta) (as defined in Floc'hlay et al., 2023). (**B-G**) UMAP plots of *phtm*, *sro*, *ImpE1*, *ImpL2* and *ImpL3* mRNA expression in uninjured (left) vs. injured (right) larval wing discs. Dotted circles highlight enrichment in alpha/beta wound populations. (**H**) Bubble plots representation of data in **B, D-G**. (**I**) IGV views of EcR Cut&Run peaks mapped using a public dataset (Uyehara and McKay, 2019).

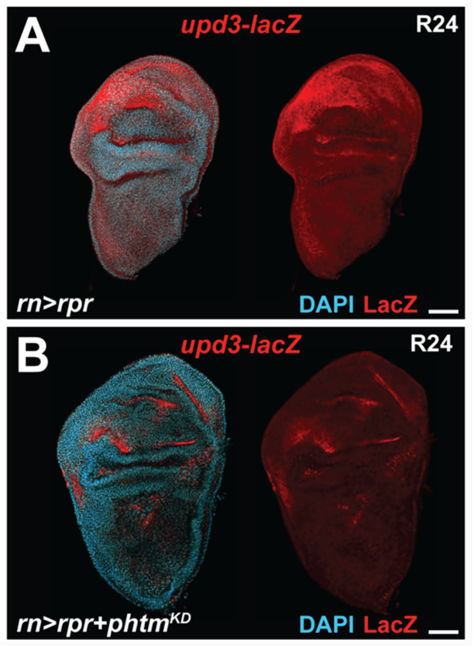


**Figure S5. Optical projection of *phtm*$^{KD}$ effect on *upd3-lacZ* expression in *rn>rpr*$^{ts}$ injured discs.** Z-stack projection of two R24 *rn>rpr*$^{ts}$ injured discs carrying the *upd3-lacZ* reporter and co-stained with DAPI. (**A)** Injury alone, (**B**) with *phtm*$^{KD}$ included (red=anti ßgal, cyan=DAPI).

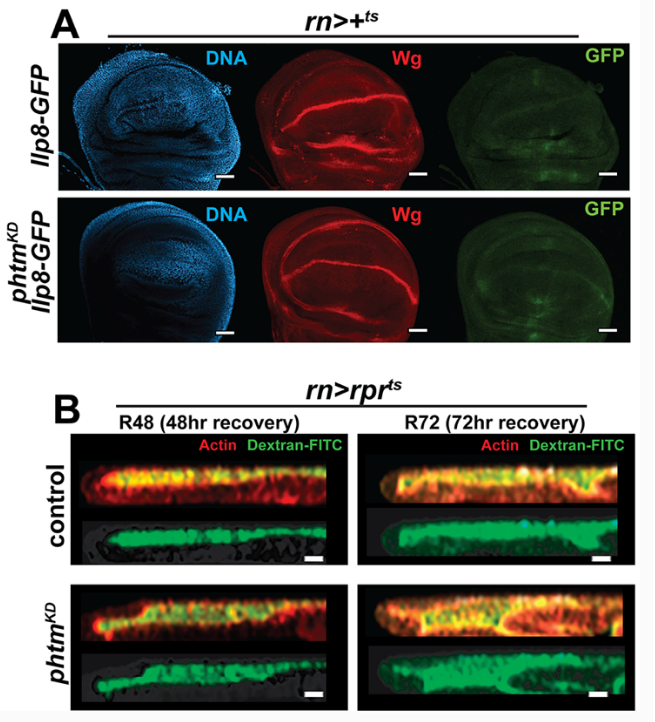


**Figure S6. *Ilp8-GFP* expression and epithelial permeability are unaffected by *pthm*$^{KD}$.**

**(A)** Representative images of *Ilp8-GFP* expression (green), DAPI (blue) and Wg (red) in mock injured (*rn*>+$^{ts}$) wing discs with and without *phtm*$^{KD}$. (**B**) Representative orthogonal views through the wing pouch of injured R48 or R72 discs +/- *phtm*$^{KD}$ incubated with fluorescein (FITC) conjugated dextran (green) and co-stained with phalloidin (F-actin; red). Phalloidin staining marks actin of peripodial and disc proper cells, which enclose a central lumen. FITC-dextran accumulates to similar levels in the lumens of injured R48 and R72 discs with or without concurrent *phtm*$^{KD}$. Scale bars=10µm.

**Table S1. Sequences for DNA primers for qPCR. Related to Figure 4.** Pairs of DNA sequences (listed 5'-to-3') used to assay expression levels of the corresponding mRNAs

| REAGENT or RESOURCE | SOURCE | IDENTIFIER |
|---|---|---|
| Primer pair for *rp49*:<br>GCTAAGCTGTCGCACAAATG<br>GTTCGATCCGTAACCGATGT | Integrated DNA Technologies | N/A |
| Primer pair for *dilp8*:<br>TGGTCATCGGAGTCTGTTGC<br>TTTTGCCGGATCCAAGTCGA | Integrated DNA Technologies | N/A |
| Primer pair for *ets21c*:<br>GTGCCAACAGAGGCCGATTA<br>CTGTTGGTGGGAACTTCCGT | Integrated DNA Technologies | N/A |
| Primer pair for *upd3*:<br>GCTGACCTTCCAGCAGAAAT<br>TGCTGTGCGTTTCGTTCA | Integrated DNA Technologies | N/A |
